# Impact of regulative noise exposure to biodiesel production due to enhanced lipid droplet production in *Saccharomyces cerevisiae*: Preliminary results from a laboratory experiment

**DOI:** 10.1101/2020.11.22.387878

**Authors:** Reetesh Kumar

**Affiliations:** Department of Biochemistry, Panjab University, Chandigarh, INDIA

## Abstract

Lipid Droplet (LD) is a ubiquitous cellular organelle that stores natural lipids as an energy and carbon source. It has emerged as a highly active organelle, engaged in lipid synthesis, protein storage, protein degradation, transportation, and metabolism. It stores natural lipids in the form of triacylglycerols (TAG) and steryl esters. TAGs consider promising biotechnological importance to produce biodiesel; thus, LD is considered a tremendous scientific concern in the modern era. The TAG accumulation is found in various feedstocks, but amongst the microorganisms becomes an evident alternative against animal and plant-derived sources due to economic reasons. Amid microorganisms, the *Saccharomyces cerevisiae* is a better alternative for industrial utilization but has low production of TAGs. Thus, to enhance the LD concentration, novel research was designed to induce alternate high and low sound frequency at a regular interval on a yeast model organism. The control and treated yeast samples further investigated using biochemical, biophysical, and computational tools to conclude that cells increase lipid droplet production under regulative noise exposure. The results endorsed that noise induces yeast LD yield is significantly higher than control, which could be considered a milestone in the biodiesel industry development and the biodiesel policy. This analysis also helps researchers to understand the novel function of LDs and their regulation in cell metabolism.

## Introduction

Lipid Droplet (LD) consists of neutral lipids bounded by phospholipids monolayer (Bartz et al. 2007), and its size varies from 20-40 nm to 100 μm (Stobart, Stymne, and Höglund 1986). It involves physical association with Mitochondria, Endoplasmic Reticulum, and Peroxisomes (Goodman 2008; Murphy, Martin, and Parton 2009) and considered a repository for biological membrane building intracellular protein storehouse (Cermelli et al. 2006). Besides this fact, LDs also employ a pivotal role in cell division (Yang et al. 2016), rotavirus replication (Cheung et al. 2010), structural and core protein assembly of dengue (Samsa et al. 2009), and hepatitis-C virus (Colpitts et al. 2015; Filipe and McLauchlan 2015; Miyanari et al. 2007). The interaction of viral protein associated with proteins linked with LDs has also been studied in numerous positive-strand RNA viruses (Cheung et al. 2010; Coffey et al. 2006; Filipe and McLauchlan 2015; Lyn et al. 2013; Samsa et al. 2009; Villareal et al. 2015). More recently, LDs accumulation increased in SARS-CoV2 patients, which act as fuel for SARS-CoV2 replication and consider as drug candidates to cure this pandemic virus (da Silva Gomes Dias et al. 2020). LD also regulates intracellular bacteria-fighting machines and acts as a regulative role in innate immunity in mammalian cells (Bosch et al. 2020). Thus, LD acts as an imperative role in the inflammatory process and infection pathogenesis (Herker and Ott 2012; Pereira-Dutra et al. 2019). On the other hand, LDs growth implies an accumulation of TAG to its core; and it notifies that TAG synthesis control by LD growth. Since TAGs might process through the transesterification produce industrial biodiesel (Atadashi et al. 2012); thus, LDs can use as a precursor for biodiesel-based factories.

Biodiesel is a type of renewable energy source formed from lipid (oil) derived from animal oil/fats, vegetable oil, tallow, and waste oils (Alptekin, Canakci, and Sanli 2014). It is considered eco-friendly, non-toxic, and pondered as a better alternate solution for fossil fuel, which produces various harmful matters, such as carbon monoxide, nitrogen oxide, and un-burnt hydrocarbon smoke, etc. (Singaram 2009). Under the European Academies Science Advisory Council (EASAC), biodiesel advancement classifies into four generations (Aro 2016). The first-generation biodiesels produce from edible feedstock such as Soybean oil, Palm oil, Olive oil, etc. (Meneghetti et al. 2007). In contrast, the non-edible feedstock is considered second-generation biodiesels (Bhuiya et al. 2014). Moreover, the third-generation biodiesel could produce from a microorganism that provides elevated growth and productivity (Carere et al. 2008). In contrast, the fourth generation production is from genetically modified algae (Lü, Sheahan, and Fu 2011). Albeit, feedstock selection is the major problem, where the use of animal and plant feedstock for biodiesel production is directly competing with human food and land resources. Thus, to oppose this challenge, oil-accumulating microbe such as bacteria, microalgae, yeasts, and other fungi have been broadly used as a secret cell factory stage for upcoming bio-refineries.

Moreover, among first and second-generation feedstock, palm oil produced maximum oil yield (>5000 kg/Ha/year) but still lacking much behind with third-generation yeast yield (1.38 L/m2/day) (Niehus et al. 2018). Therefore, altogether preferring yeast over other microbes to restore fatty acids for biodiesel production is a significant concern currently. In equivalence, the yeast genomes genetic manipulation has also increased lipid content, such as triacylglycerol (TAG) and starch, that can produce several distinct biofuels, biodiesel, and ethanol (Lin et al. 2013). Nevertheless, the overproduction of LDs in the wild yeast is still a matter of current research consider the economic point. In such a case, the present study has designed to execute LD production using noise as a tool on eukaryotic cellular systems using yeast as a model organism.

Though, the impact of noise on animals (Algers, Ekesbo, and Strömberg 1978; Castelhano-Carlos and Baumans 2009; Kight and Swaddle 2011), plants (Ghosh et al. 2016) as well as on microorganisms (Sarvaiya and Kothari 2015) already been studied based on its absorption, production, and transmission. However, not much consideration has compensated for biological effects induced by audible noise, as reported in the case of *E. coli* (Shaobin et al. 2010). Therefore, in the present study, the audible sound stimuli generation device is forced to alternately generates high and low-frequency sonic vibration from a speaker or resonance fork. The intersecting part of the experiment is that LD production increases without changing the reserved yeast’s genetic makeup. Thus, this procedure could increase the number of lipids with minimum expenditure and the shortest possible time. In this view, the study will be a stepping stone in lipid droplet-based research, not because it reveals the condition-specific alteration in lipid biogenesis but also because it increases the knowledge about cellular pathways involved in homeostasis and cell health. Similar genomic and proteomic analysis correlated with the higher eukaryotic organism. (e.g. human) can be extended from the information of yeast LD proteome. Therefore, the implication for human studies may help in the understanding of the metabolic disorder, cancer neurodegenerative disorders and more recently on COVID-19 studies.

## Result and Discussion

The constant noise/sound frequency already employed in the yeast culture for metabolic analysis (Aggio, Obolonkin, and Villas-Bôas 2012) directs various biotransformation responses. The experiment reveals under constant higher, and constant lower frequency, the same/or different metabolites expressed, which conclude that the noise/sound frequency could be used as potent manipulating cell metabolism and proliferation tools in future research based on yeast model organisms. Moreover, noise/sound affects protein expression by managing the transcription, translation, and degradation rate already used as a tool for expression analysis (Liu, François, and Capp 2016; Mundt et al. 2018). The noise expression in the target gene/s (TGs) results from fluctuations in transcription factors (TFs), a phenomenon called noise propagation. A small variation near the critical TF threshold can initiate an expression switch of TG and alter cellular phenotypes (Hooshangi, Thiberge, and Weiss 2005; Tkacik, Callan, and Bialek 2008). Thus, noise results from randomness, but some genes are steadily noisier than others (Lehner 2008; Newman et al. 2006). Thus their positive and negative selection recommend using a developable characteristic tool for expression analysis.

As such, noise/sound induces the yeast cell, which controls the cell size and could use as phenotypic sampling as already analyzed (Liu, François, and Capp 2016). Molecular noise was also implemented on yeast cells to create and organize cell cycle variability monitoring by G1 cyclin using statistical analysis (Di Talia et al. 2007), which endorsed that the daughter cell shows more robust size control than mother cells. On the other hand, Ime1p, an essential regulator of meiosis in the yeast cell, shows active variability in the sporulation pathway under noise exposure (Nachman, Regev, and Ramanathan 2007). Besides, noise-induced green fluorescence protein (GFP) expression levels during transcriptional and translational conditions were analyzed using yeast as a model organism (Blake et al. 2003), which deduced the maximum expression under intermediate noise conditions. Therefore, an alternate high and low frequency was used in this experiment to get a better picture of protein expression. As noise/sound selection, select the enchanting word “AUM” because; its intensity increases and decreases gradually makes an intermediate sound exposure, where protein expression was maximum as reported (Fraser et al. 2004). The audio file of noise “AUM” reports in the Supplementary audio file SA1. Another reason is that it is one of the shortest audible sound and less chance of error in consecutive experiments, and easy to manage in repeated experiments. The intensity of noise was 100 Hertz as lower frequency and 10,000 Hertz as higher frequency, respectively, which already consider under constant frequency-based metronomic analysis on the yeast (Aggio, Obolonkin, and Villas-Bôas 2012). In such a case, under the alternate high and low sound frequency rate of glycolysis and TCA cycle found an increased rate, which under regulative metabolism covert to LDs.

Though the LD considered as fat storage, in the last 20 years, which pondered as dynamic organelles that are involved in lipid metabolism, cell signaling, inflammation, membrane biosynthesis, neurodegenerative disorder, biodiesel, and cancer (Farese and Walther 2009; Fujimoto and Parton 2011; Welte 2015). LDs also play an essential role in embryo development to maintain histone protein balance (Li et al. 2012; Tatsumi et al. 2018). It also involves in metabolic diseases such as atherosclerosis (Faber et al. 2001; Paul et al. 2017), viral replication (Miyanari et al. 2007), diabetes (Olofsson et al. 2011), and might adjust the rate of sterol biosynthesis by inhibiting Erg1 (Leber et al. 1998). Besides, LDs are also involved in protein trafficking and their maturation within the cell, but still do not know the process involved independently or are symptomatic of the general cellular protein trafficking pathway (Bersuker and Olzmann 2017). Altogether, many questions need to be solved, which could be considered a promising target for future research. Though, amongst the LDs, it directly involves biodiesel production (Tsai et al. 2015); hence LD enrichment is a prerequisite for future research to oppose the energy crises, which is the principal objective of this experiment. In such quest, among all available feedstock for LDs production, the yeast gives the impression that the most adapted organism accumulates lipid predominantly in TAG (Steen et al. 2010). Another benefit is that the yeast process’s qualitative and quantitative output is much advanced to plant cultivation using low-cost material, short life cycle, less labor requirement, and less care for the location and climate. Thus, yeast is an immense opportunity to utilize as a starting material to restore fatty acids for biodiesel production through transesterification. Soybean and rapeseed oil is the most common feedstock for biodiesel, which contains a similar fatty acid profile as yeast (Beopoulos and Nicaud 2012), which assists yeast as a suitable candidate for biodiesel production. Hence, the experiment organized using noise/sound exposure, which will develop new challenges for cost-effective biodiesel production using *S. cerevisiae* established factories.

Thus, the experiment design, where the natural yeast strain of *S. cerevisiae* was cultured in YPDA media using glucose as the mere carbon source. The culture was further inoculated at 28°C and separated into two identical clones as control and treated flasks, using the shaker incubator. The alternate high and low audible sound frequency waves were applied to the treated flask, whereas the control flask was grown in silent conditions. The exposure was given, especially in the experimental block under 8 hours, which does not physically contact yeast cells, subsequently induces only cellular metabolism. Thus, noise exposure bypasses contamination in the yeast culture, which could occur due to drug incubation and bacterial contamination. Finally, total protein from control and noise treated cells were extracted and quantified (Kushnirov 2000). A further similar amount of proteins was separated by 2-DE using 3±10 pH range IEF strips to generate a the SDS gel spot. The segregated spot among control and treated gels observed along with the range of molecular weight (Mr) and Isoelectric points (PI), as shown in Fig. 1A. Under highly basic conditions, fewer spots measured; however, spots produced expressively more abundant in the 35 to 65 kDa and 3 to 7 pI range. Melanie9 software (Berth et al. 2007) used further to detect the specific spot intensity three-dimensionally (Supplementary Fig. SF1) and judge precisely the major difference in their spot intensities. Finally, 14 spots were selected and subjected to MALDI-TOF/TOF analysis for their identifications. The proteins were chosen based on the MASCOT score and top identified peptides using the m/z ratio, as shown in Table 1. Out of 14, 13 spots were up-regulated, and one spot was down-regulated under noise treatment, as shown in Fig. 1B. The identified up-regulated spots were TDH2 (Glyceraldehyde-3-phosphate dehydrogenase, isozyme 2), TDH3 (Glyceraldehyde-3-phosphate dehydrogenase, isozyme 3), GPM1 (Tetrameric Phosphoglycerate Mutase), ADH1 (Alcohol dehydrogenase), ENO2 (Enolase II), PGK1 (3-phosphoglycerate kinase), HSP60 (heat shock protein), TEF1 (Translational elongation factor EF-1 alpha), TSR3 (20S rRNA accumulation protein 3), ARP2 (Actin-related protein 2), IML3 (Increased minichromosome loss protein 3), MTC2 (Maintenance of telomere capping protein 2), COX5A (Cytochrome c oxidase polypeptide 5A) and the down-regulated spot was DYL1 (Dynein light chain 1). Further, a protein-protein interaction network (PPI) was constructed by STRING, a Cytoscape plug-in to improve the functions and interactions of recognized proteins. The protein interaction network analyzes with a cut-off confidence score of 0.40 as default support that selected proteins have strong co-expression and co-occurrence in the cellular environment (Fig. 1C). The predicted protein-protein interaction evaluation as analyzed by STRING, endorse that the most potent interaction found between TDH3-ENO2, TDH3-PGK1, TDH2-ENO2, TDH2-PGK1, PGK1-ENO2, PGK1-GPM1, GPM1-ENO2, and ENO2-TDH3 (Combined association score: 0.999) and lowest interaction among HSP60 and ADH1 (Combined association score: 0.401) as shown in Supplementary Table ST1. The MCODE, another Cytoscape plug-in used on STRING data to find a densely connected region based on network-based topology, as shown in Fig. 1D. MCODE suggested cluster contains HSP60, TEF1, ADH1, PGK1, GPM1, ENO2, TDH3, and TDH2, as shown yellow color, linked with COX5A as blue. The arrow thicknesses illustrate the level of strength in their interactions. MCODE suggested top score depicts the significant interactions involved in energy derivation by oxidation of organic compounds, ATP metabolic process, ATP catabolic process, nucleoside phosphate metabolic process, glycolysis, and fermentation, and ethanol biosynthetic process involved in glucose fermentation to ethanol as sown in Supplementary Table ST2. The expressed protein under Wikipathway illustrates among these proteins, TDH2, TDH3, PGK1, GPM1, and ENO2 are directly involved in Glycolysis and Gluconeogenesis. However, ADH1 involve in transport between cytosol and mitochondria, which play an essential role in the conversion of acetaldehyde to ethanol (Supplementary Fig. SF2). In brief, ARP2, COX5A, ENO2, GPM1, HSP60, PGK1, TDH2, TDH3, and TEF1 were share mitochondria as a cellular compartment as shown in Supplementary Table ST3, where ARP2 play a direct role in the actin cytoskeleton, which gives strength to the cell (Veltman and Insall 2010). On the other hand, COX5A is a Cytochrome C oxidase of 17 kDa found in the inner mitochondrial membrane, which executes the role in ATP synthesis as the last enzyme in the mitochondrial electron transport chain (Hartley et al. 2019). HSP60, another induced protein act as a mitochondrial chaperon, helps fold and maintain around 30% cellular proteins (Ranford, Coates, and Henderson 2000), helping other proteins to bring in the mitochondrial matrix (Koll et al. 1992). Thus, the result informed that under noise exposure, the treated cells increase their metabolism during glycolysis, and eventually, the TCA cycle starts in mitochondria. Due to sufficient glucose, acetyl-CoA is formed in the mitochondria matrix mostly by oxidative decarboxylation of pyruvate (from glucose, amino acids). The acetyl-CoA is unable to go through the mitochondrial membrane, and thus it is transported in the form of citrate when citrate is not necessary for the TCA cycle. Subsequently, the TCA cycle’s down-regulation accumulates citrate in mitochondria and distributes surplus citrate from mitochondria to the cytosol. In the next cycle, ATP-citrate lyase (ACL) cleaved citric acid into cytosolic acetyl-CoA (AcCoA) and OAA (oxaloacetate). Acetyl CoA does not have sufficient energy to enter the fatty acid synthesis; therefore, acetyl-CoA carboxylase (ACC) converts the Acetyl CoA to Malonyl CoA (MalCoA). Further, the FA synthase complex (FAS1 and FAS2) acts on MalCoA to construct 16 carbons Acyl-CoA, which transport into the Endoplasmic Reticulum (ER) for elongation (Farese and Walther 2009). The elongation process depends on ATP and NADPH’s availability for the construction of Fatty acid (FA). Eventually, three FAs condenses with glycerol to form one molecule of Triacylglycerol (TAG). In such a case, ACC activity was analyzed under control and treated conditions, as shown in Fig. 2A. The result revealed that under-treated conditions, the ACC concentration was higher than control cells, which execute a similar result, as shown in the oleaginous yeast *Yarrowia lipolytica* (Tai and Stephanopoulos 2013). TAGs are also produced by DAG acyltransferase, which uses Acyl-CoA as acyl donor (Dahlqvist et al. 2000). The reactions located on the surface of ER and the LD, which formed due to the presence of TAGs and most of the relevant enzymes over there.

**Figure 1.**
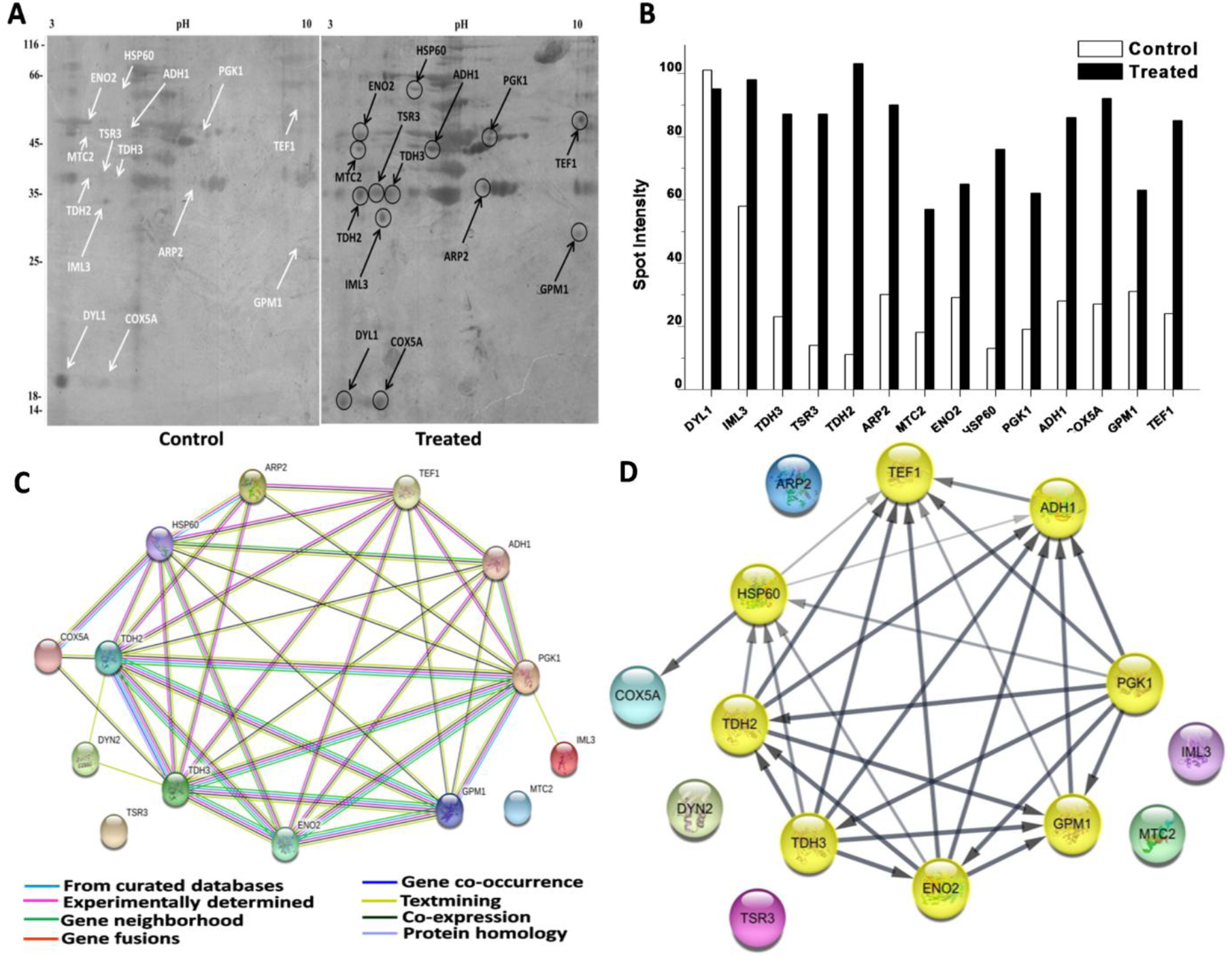
(A) Two-dimensional electrophoresis (2-DE) separation of control and treated proteins. An equal amount of both samples loaded for each IEF strips (pH 3-10) followed by the second dimension on 12.5 % SDS-PAGE. Proteins exposed by silver staining. The relative molecular weight marker used to estimate the molecular weight of concerning proteins shown extreme left. (B) Spot Intensity of expressed proteins measured by Melanie software. White and Black columns show control and treated expression, respectively. (C) Analysis of protein-protein interaction (PPI) network by String, a Cytoscape plug-in. PPI is accessible for noise expressed proteins under a confidence level of 4.0 as default. Different lines illustrate the type of evidence used in predicting the association between expressed proteins. The concerned edges and lines represented, as mentioned below, that predicting the association of the protein. (D) String software predicted interaction analyzed using MCODE, a Cytoscape plug-in. The MCODE module was detected primary cluster (Score; 7.714, nodes; 8 and edges; 27) as shown yellow. Line thickness indicated the strength of confidence, and arrow orientation illustrates their interactions.

**Table 1.**
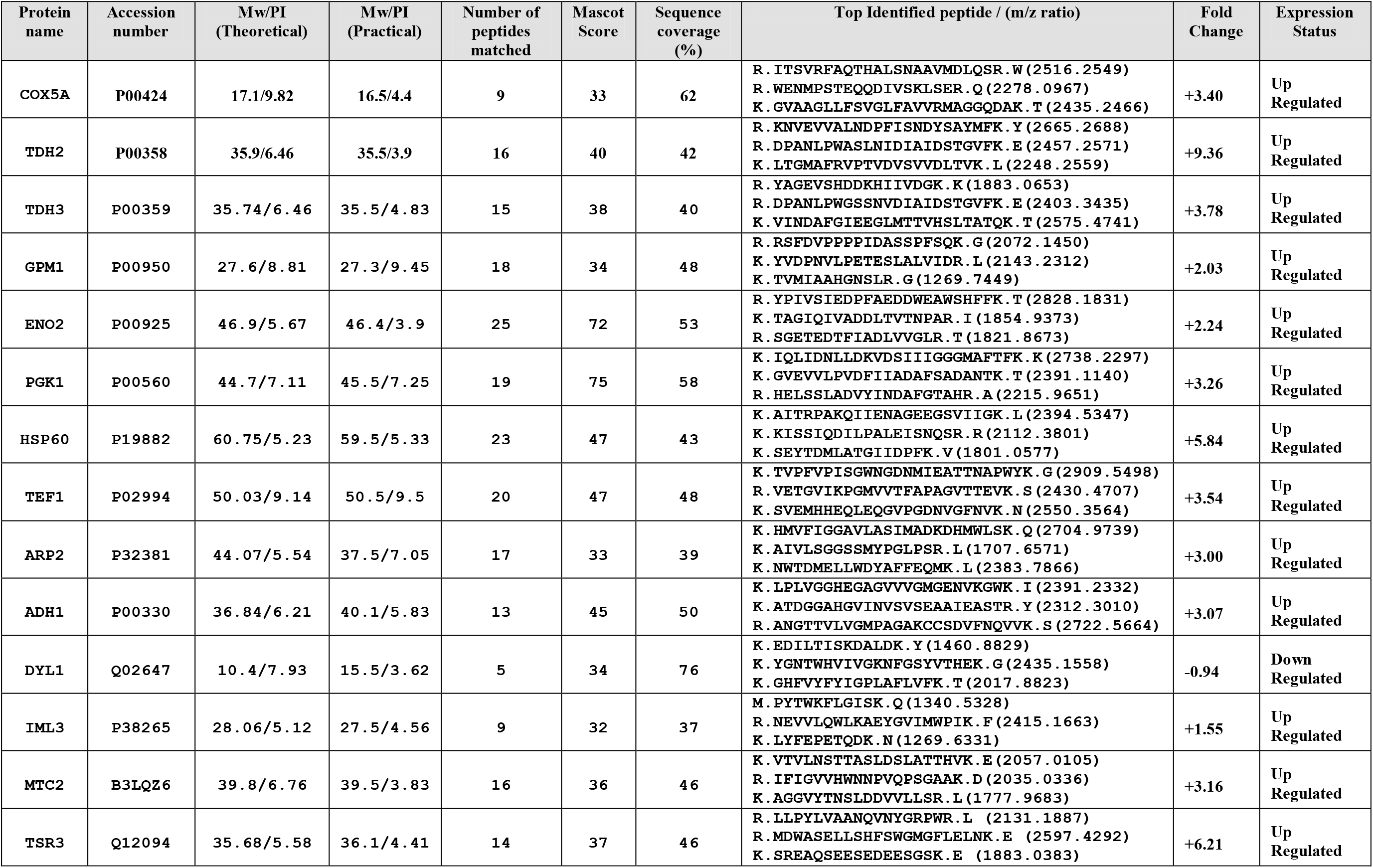
Differentially expressed proteins in yeast culture executed under noise treatment identified using MALDI-ToF/ToF mass spectroscopy.

**Figure 2.**
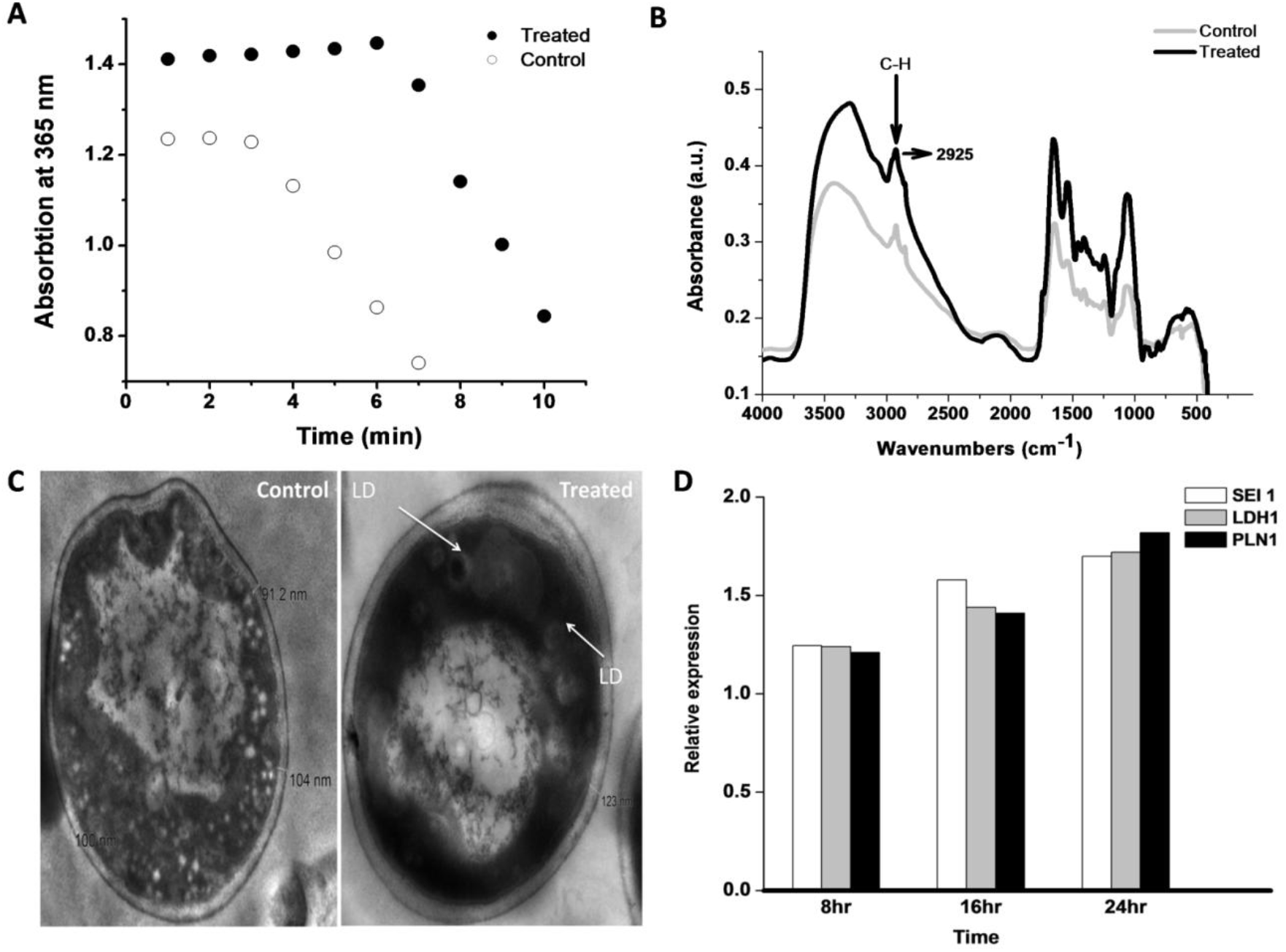
(A) Acetyl-coenzyme A carboxylase (ACC) assay: Spectrophotometric assay of control and treated culture to determine the ACC activity. Malonyl—CoA formed by acetyl-CoA carboxylation, which reduced to 3-hydroxypropionate using 2NAPDH, was observed spectrophotometrically at 365 nm. (B) FTIR spectra of yeast cells under control and treated samples. (C) Transmission Electron Microscopy (TEM) image under control and treated condition. (D) Real-time PCR Real-time PCR measurement of Seipin1(SEI1), Lipid Droplet Hydrolase1 (LDH1), and Perilipin-1(PLN1) expression undertreated condition. The experiment was executed under 8 hr, 12hr, and 24 hr incubation duration.

LDs consist of high fatty acid content; therefore, FTIR was conducted under control and treated cells, as shown in Fig. 2B. The result reports that at 2925 wave number treated peaks spectra was 30% higher than the control sample. The 2925 wave number is specific for C-H bond stretching, representing the strong indicator of fatty acid present in the treated sample (Forfang et al. 2017; Mihoubi et al. 2017). The result also illustrates that new fatty acid form under noise exposure was absent in the control condition. Further, to execute the LD morphologically at the cellular level, TEM analysis was done (Radulovic et al. 2013). The cell membrane thickness under control (104 nm) and treated condition (123 nm) shown in Fig. 2C, support that treated cell protects them from noise/sound-induced condition and subsequently promote more glycolysis and store energy in the form of LDs. LD-specific protein Seipin-1 (H. Wang et al. 2016), Lipid Droplet Hydrolase 1 (Thoms et al. 2011), and Perilipin-1 (Gao et al. 2017) also used to detect Real-time PCR analysis under control and induced condition, which illustrated in Fig. 2D. Among them, Sepin1 (SEI1) is the specific protein for lipid droplet maturation (Sui et al. 2018; Zoni et al. 2019), which function as membrane anchors to facilitate LD formation and support their growth. Another LD-specific protein, Lipid Droplet Hydrolase1 (LDH1) used in lipid homeostasis and mobilization of LD (Debelyy et al. 2011), containing a characteristic catalytic triad (GxSxG motif), whereas, Perilipin (PLN1) implicated in the stability of LDs and consider as the doorkeeper of lipolysis (Sztalryd and Brasaemle 2017). The LDH1, PLN1, and SEI1 proteins under 8 hr, 12hr, and 24 hr incubation increase their expression, reports to LD concentration enhanced under induced condition. LD enrichment application in cell viability was analyzed by spotting test under serial dilution, as shown in Fig 3A. The control and treated cells were first taken after 3 hours of experiment, and test samples were taken at every 9th-hour duration, which clears to show that treated cells survive a bit more time than control conditions. As analyzed, the up-regulated proteins found in the cytoplasm and mitochondria; hence, mitochondria was confirmed using MitoTracker Deep Red FM fluorophore. The control and treated samples subjected to Mitotracker dye and the stained cells were analyzed morphologically by confocal microscopy, as shown in Fig 3B. The result illustrates that in undertreated condition, cells were more stained in contrast to control cells. In support of mitotracker dye, LD specific fluorophore Nile red (Greenspan and Fowler 1985) also analyzed by confocal microscopy. The Nile red is particular for intracellular lipid droplets described by Greenspan *et al*. (Greenspan, Mayer, and Fowler 1985), which nullifies the presence of lipid-water interface undertreated condition. The LD analyzed using a confocal microscope, as shown in Fig 3C, depicts that treated cells were more stained compared to control cells. LDs presence under control and treated condition were also analyzed by Fluorescence microscopy using a TRITC filter set, which illustrates that cell under-treated condition more flourished, as shown in Fig. 4. Further, the stained cells (using mitotracker and Nile red) under control and treaded condition counted by flow cytometry. The flow cytometry results were analyzed using FlowJo software (Chen et al. 2015), which selected forward-scattered light (FSC) vs. side-scattered light (SSC) as an axis parameter and formed a scatter plot. The results illustrate that control LDs cells were 33559 in number compared to 65173 treated LDs cells using almost similar unstained cells, as shown in Fig. 5A. The increase in 94% of cells under-treated LDs with 892 mean values than 189 means of control LDs inform that LDs concentration increases under treatment. The mitotracker stained cells also evaluated as shown in Fig. 5B, where treated cells were more than 2 times higher than control cells with 4 times higher mean value, which undoubtedly informs that LD production increases due to the distribution of surplus citrate from mitochondria. Further, the 48th-hour samples tested for cellular ROS level using Dihydrorhodamine 123 dye. The result discloses that under the treated condition, the ROS mean level was 866 compared to control 1319, with almost similar stained cells under both conditions, as shown in Fig. 5C. The lower the mean value under-treated condition illustrates that the cell may decrease the intracellular ROS level during LD production. LD acts as an antioxidant in Drosophila’s stem cells recently studied (Bailey et al. 2015), supporting this analysis. As ROS level decrease undertreated condition, the cell proliferation assay was also executed. In such a case, CFSE dye was used for flow cytometry followed by FlowJo software for analysis as shown in Fig. 5D. The result informs that under each proliferation steps of treated cells, CFSE cell count was high in number in comparison with control CFSE cell count. Cell proliferation and lipid metabolism as already discussed using epidermal growth factor (EGF) signalling pathways (Eling and Glasgow 1994), which inform that linoleic acid metabolites control the EGF pathway and help in cell proliferation. This analysis clear to illustrate that cells under treated condition were decrease ROS level and proliferate in a more smooth mode in contrast to the control condition. Thus, cells under-treated conditions were more proliferate and excess energy stored in the form of LDs. Consequently, this study first time explore that noise could play a very important and direct role in LD accumulation and considered as a stepping stone in the field of LDs based research.

**Figure 3.**
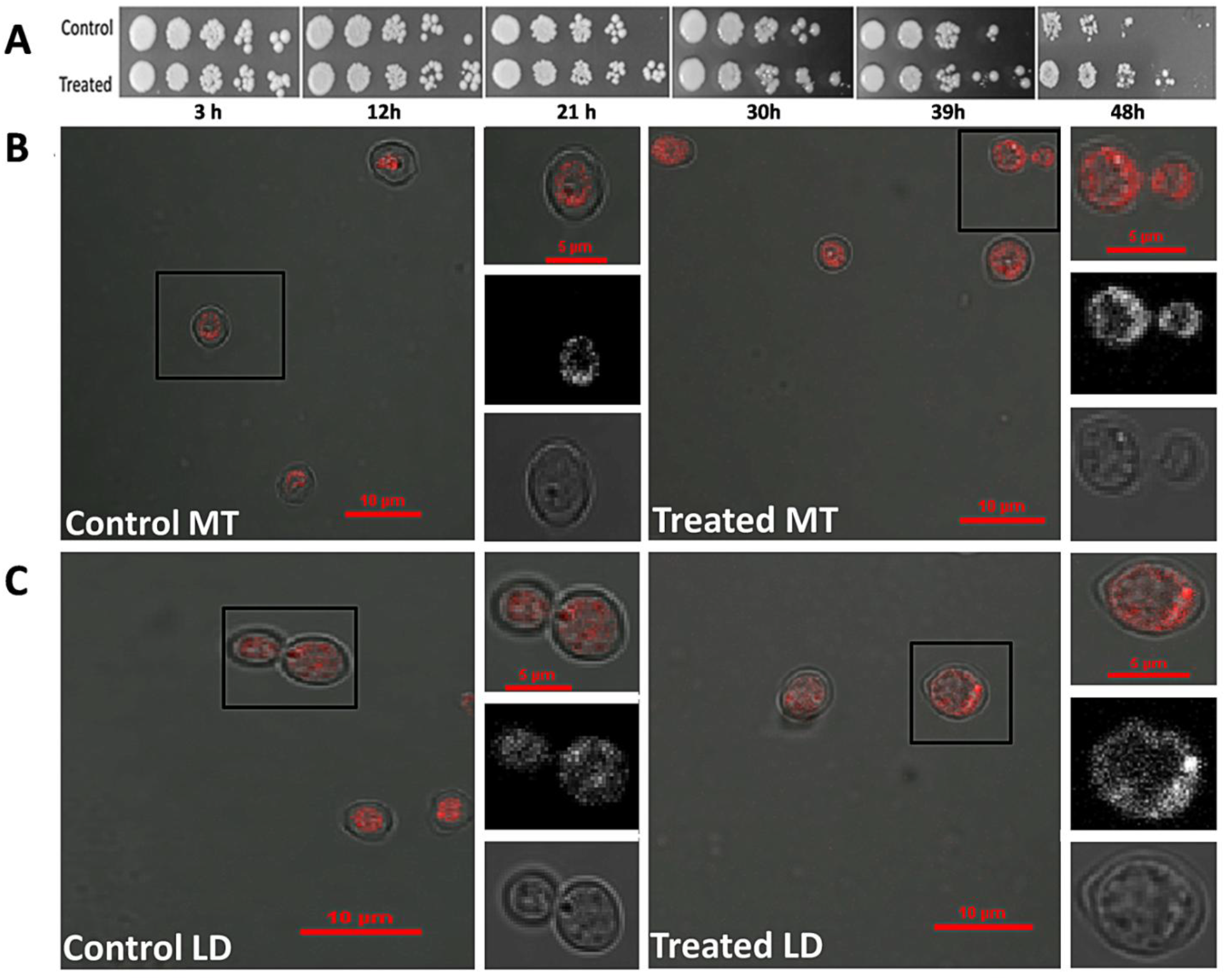
(A) Spot test analysis: Equal numbers of yeast cells were spotted on media under control and treated conditions. The comparative numbers of yeast cells were spotted on the plates shown at regular intervals from the experiment. (B) Confocal microscopy of yeast cells using MitoTracker (MT) deep red stain under control and treated conditions. Inset illustrates the selected cells under different visual conditions. Scale bar included. (C) Confocal microscopy of yeast cells using Nile red as intracellular LD specific dye under control and treated conditions. Inset illustrates the selected cells under different visual conditions. Scale bar included.

**Figure 4.**
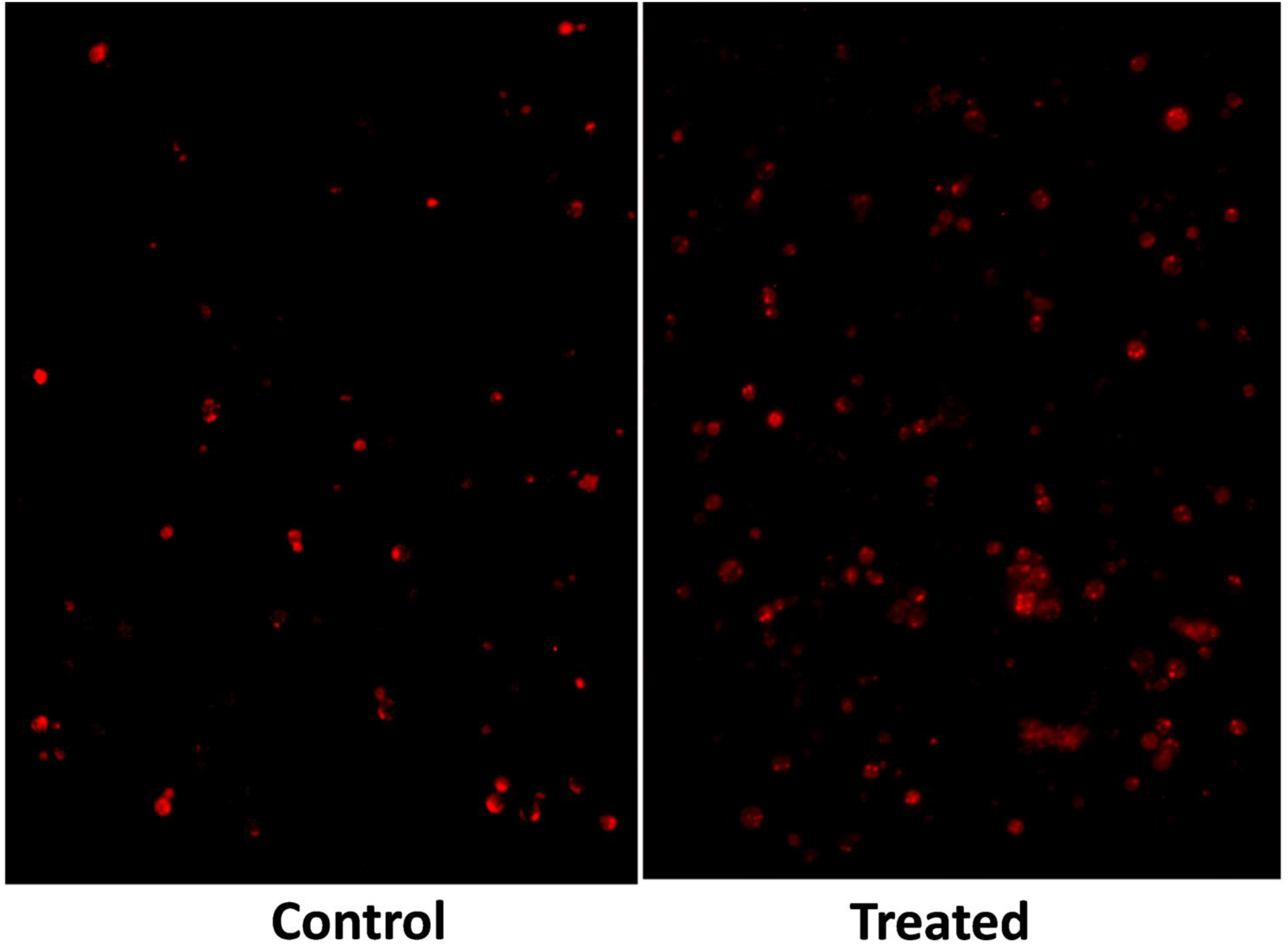
Fluorescence microscopy analyses of Nile red-stained LDs in yeast cells under control and treated condition.

**Figure 5.**
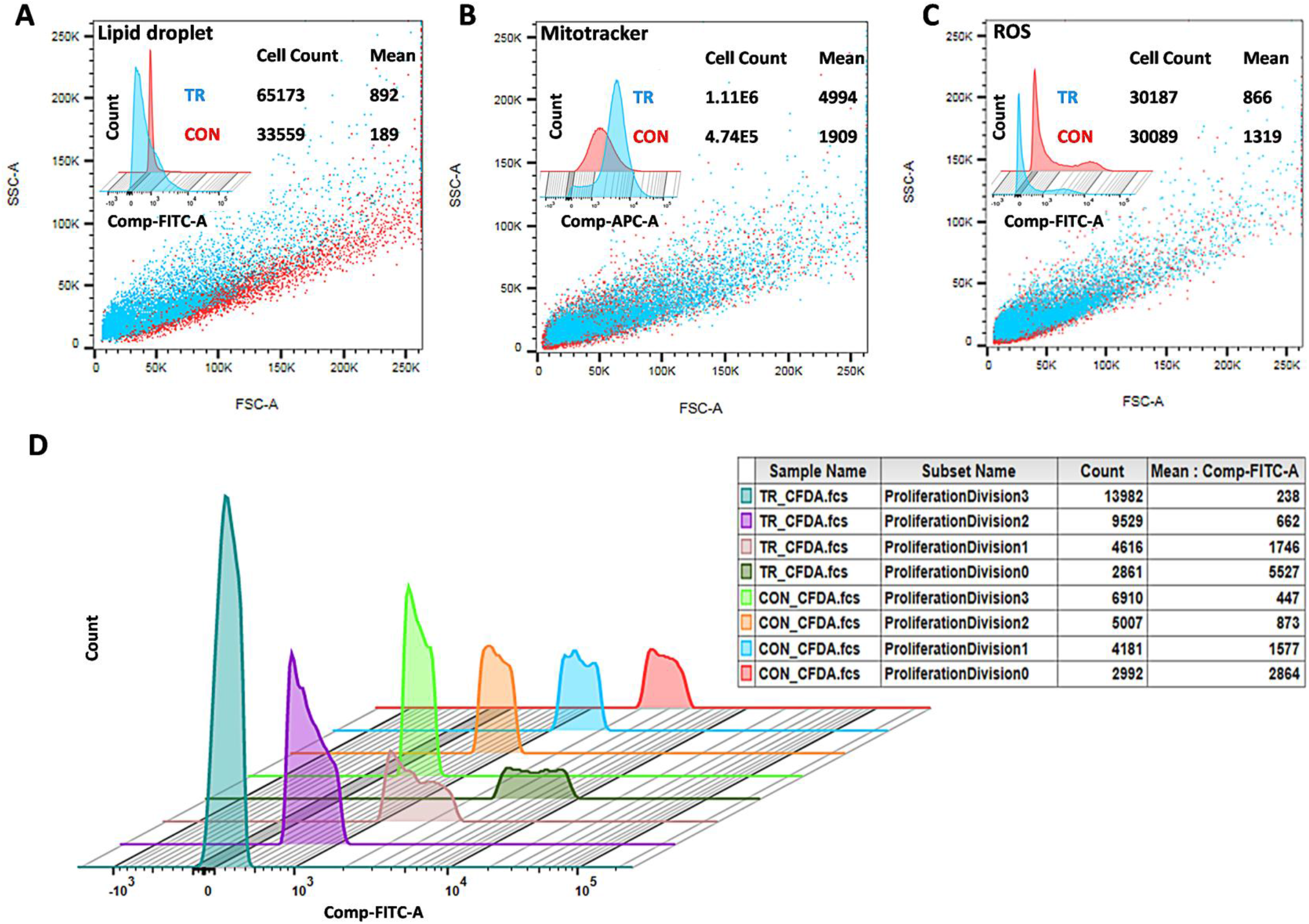
(A) LD detection using FACS under control and treated condition. (B) Mitochondria detection using FACS under control and treated the condition. (C) ROS detection using FACS under control and treated the condition. SSC vs. FSC plot of control and treated sample showed red and cyan in color. Cell count and mean value mentioned right upper corner. Overlaying is a powerful visual tool for cytometric analyses shown upper left corner. (D) Histogram Overlay using cell proliferation dye CFSE under control and treated conditions. The table depicts the sample name under different proliferation division with cell count and their mean values. Control is illustrated as CON and treated as TR under every condition.

## Conclusion

2D-PAGE, MALDI, Gene Ontology illustrates that noise exposure activates the yeast cell’s lipid droplet production. Further, String and MCODE inform that the noise exposure motivates the cell to increase ATP metabolism during glycolysis. Due to noise exposure, no contamination will increase in the cell, which is the major strength of this work. Additionally, TEM analysis exposes that cell wall thickness increases, which informs that cells want to increase their size due to more ATP production. Subsequently, LD concentration increases accordingly to maintain the surplus of energy. LDs conformation check by LD detection kit, FTIR. Cellular ROS levels decrease during noise exposure analyzed by Dihydrorhodamine 123 dye. Cell proliferation also analyzes by CFSE dye, which informs that cells under noise exposure proliferate more rapidly. Altogether, protein expression analyzes using bioinformatics Cytoscape tools, which inform that LD production occurs during frequent noise exposure. Thus, this is the simplest way for LD production using yeast as a model organism. LD use as a precursor in biodiesel based factories, so this work would be meaningful and robust for LD production.

## Acknowledgments

The author is thankful to Dr. Robin Joshi from the Institute of Himalayan Bioresource Technology, Palampur (IHBT), Himachal Pradesh, INDIA for 2-DE, and MALDI experiments. The author acknowledges Mrs. Bhupinder Kaur from the Post Graduate Institute of Medical Education & Research (PGIMER), Chandigarh, INDIA, for Flow Cytometry experiments. The author, thanks to Mr. Deepak Bhatt from the Institute of Microbial Technology and Post Graduate Institute of Medical Education & Research (Imtech), Chandigarh, INDIA for Confocal Microscopy. The author is thankful to Dr. Charan Singh Rayat from PGIMER for Transmission electron microscopy (TEM) experiments. The author thanks Dr. Sukesh Chander Sharma, Department of Biochemistry, Panjab University, Chandigarh, INDIA, for supporting this work. The author thanks Dr. D S Kothari for Funding support from UGC.

## Declartioan of Intrests

The author declares no competing interests.

## Materials and Methods

### Culture and growth conditions

The natural yeast (*Saccharomyces cerevisiae*) used in this study purchased from Kothari Fermentation and Biotech Limited, New Delhi, INDIA. The executed culture was grown in the YPDA media (1% Yeast extract, 2% Peptone, 2% Dextrose, and 2% Agar). The inoculums prepare by transferring a single colony from the YPDA plate into 50 ml of YPD broth. Which further incubated at 30°C overnight at 160 rpm. The final cell density equivalent to 0.05 OD600 of inoculums was transferred to the duplicate 100 ml YPD broth in a 500 ml Erlenmeyer flask, and culture again incubated at 30°C at 160 rpm. Subsequently, after getting the mid-log phase (16 hours), one was transferred into the noise chamber, and another under silent condition use as a control. The noise chamber was 54 x 58 x 60 cm in dimension. A speaker delivering noise “AUM” was placed in the chamber near the inoculated flask for 8 hours at a distance of 20 cm. The noise’s intensity was measured (100-10,000 Hertz) with an Acd machine (Sarvaiya and Kothari 2015). The noise chamber was entirely covered with a glass lid as noise-proof packing to prevent any possible leakage, protection, and protection from external noise. The control inoculums place in another similar dimension chamber without speakers; subsequently, no noise generate.

### 2D Gel Electrophoresis

Control and noise treated *S. cerevisiae* cells were harvested by centrifugation at 6000 rpm for 5 min, and the pellet was washed two times with distilled water. The stock samples further prepare from the pallet according to the method described by Kushnirov (Kushnirov 2000), which subsequently performed 2-DE using a 7 cm immobilized pH gradient (IPG) strips (pH range of 3—10, Ready strip, Bio-Rad, USA).

The control and treated samples were kept in 10x ice-chilled acetone at −20°C for 3h for protein precipitation, afterward centrifuged at 7500 rpm for 10 min at 4°C to get a pellet. The desired pellet was further dried for 5 min then dissolved in freshly prepared lysis buffer (0.5% Triton X-100, 1mM EDTA, 50mM DTT, 20 mMTris (pH 7.5), 2 M Thiourea, 7 M Urea, 4% CHAPS) and processed for isoelectric focusing (IEF). Further, the strips loaded on 12% polyacrylamide SDS-PAGE for the second dimension gel system. Gels were further kept overnight in 40% methanol with 1% glacial acetic acid solution and stained using a modified silver staining method (Yan et al. 2000). Two-dimensional gel images developed using a GS-800 calibrated densitometer scanner (Bio-Rad). Three replicates of every gel were analyzed. Spot quantification accomplished using Melanie 9 software (Appel et al. 1997) employing a difference in spot intensity by aligning the spots within concerned gels and quantify the selected spots accordingly. Every spot on the master gel is considered based on presence in at least two of the three gels. The spot intensity vs. volume reflects a significantly up-regulated and down-regulated expression. The mean of differently expressed spots was calculated, and the concerned value is used as spot quantity on the standard gel. Fold change (+/-) considered the relative volume ratio (spot intensities) in the treated control gel. The molecular mass of each protein calculates by comparing it with a standard molecular marker (Bio-Rad). The isoelectric point (pI) verify by the spot positions along the pH gradient strips.

### Mass spectrometry

Differently expressed protein spots from silver-stained polyacrylamide gels were manually excised based on up-regulated/down-regulated noise treatment. The excised gel pieces destain in the mixture (1:1, v/v) of 30 mM K_3_[Fe(CN)_6_] and 100 mM Na_2_S_2_O_3_ at room temperature for 20 min. The engrossed gel pieces were vortexes until destained, washed further three times with 200 μl of Milli-Q water for 5 min, and dehydrated in 100 μl of acetonitrile. The gel samples were inflated in a 25 μl of trypsin (Sigma, USA) solution (20 μg/ml in 25 mmol/L NH_4_HCO_3_) for 30 min at 37°C and eventually incubated overnight at the same temperature. Each digested peptides extracted from the gel with 50% Trifluoroacetic acid/50% Acetonitrile at room temperature. The extracted peptide mixtures were supplemented with 0.5 μl of α-Cyano-4-hydroxycinnamic acid (Bruker) (20 mg/ml) in 0.1% trifluoroacetic acid/30% (v/v) acetonitrile (1:2) and dried at 37°C. The isolated peptides were exposed to MS using MALDI-ToF/ToF-Proteomics Analyzer (UltrafleXtremeTM mass spectrometer; BrukerDaltonics Inc. Germany). A mass standard starter kit (BrukerDaltonicsInc, Germany) and a standard tryptic BSA digest (BrukerDaltonicsInc, Germany) are used for the MS and MS/MS standardizations system. The combined MS and LIFT-MS/MS accomplished using BioTools 3.0 software (BrukerDaltonicsInc, Germany). The TOF spectra were recorded from 700 to 3500 Da in a positive ion reflector mode. Each spectrum accumulated with five hundred shots, and among them, two most abundant peptide was exposed to fragmentation study to resolve the peptide sequence. The Database accomplished using the MASCOT search engine (Version 2.1, Matrix Science, London, U.K) with Swiss Prot database (Release date, 5th May 2013; version 121; 540052 sequences). All peptide masses expected monoisotopic and [M+H+]. The other factor used for the search was as follows, enzyme, trypsin, the fixed modification, carbamidomethyl (C), variable modification, oxidation (M), parent ion mass tolerance at 100 ppm, and MS/MS mass tolerance of 0.7 daltons with one missed cleavage allowed. The identified proteins were selected based on top listed hits on the search report with extensive homology (p <0.05). The assurance in the peptide mass fingerprinting matches were based on the score level and confirmed by the matched peptides’ accurate overlapping, which shows major peaks of the mass spectrum.

### Data Processing and Bioinformatics Analysis

The identified proteins searched for their biological functions using the UniProt database (www.uniprot.org). The gene’s name was retrieved from the UniProt web server and used to construct protein-protein interactions using STRING, a Cytoscape plug-in (Szklarczyk et al. 2015). Molecular Complex Detection (MCODE), another plug-in, was used to perform network clusters based on the topology to identify the closest proteinprotein interaction networks in Cytoscape (Bader and Hogue 2003). In parallel, the pathway involved with maximum expressed protein was represent by WikiPathways, a Cytoscape plug-in (Kutmon et al. 2014).

### Fourier Transform Infrared Radiation (FTIR) analysis

FTIR analysis executed on a PerkinElmer Spectrum GX spectrometer (PerkinElmer, altham, Massachusetts, USA) equipped with a liquid N2-refrigerated MCT (Mercury Cadmium Telluride) detector. The measurement records between 4000 and 450 cm-1. Data analyzes of IR spectra performed using OMNIC software (Thermo Scientific).

### Transmission Electron Microscopy (TEM) analyses

TEM was used for intracellular changes in the morphology of organelles under control and treated yeast cells. TEM was processed, where samples were collected and fixed with 2.5% EM grade glutaraldehyde in 0.1 M sodium cacodylate buffer (pH 7.4) at 4°C. Fixed cells were placed further in 2% osmium tetroxide in 0.1 M sodium cacodylate buffer (pH 7.4). Afterward, dehydrate the cells in graded series of ethyl alcohol and be implanted in Durcupan resin. Ultrathin sections (thickness, 80 nm) of cells were cut with a Reichert-Jung Ultracut Eultramicrotome and examined using a JEM-1400 electron microscope.

### qRT-PCR

Total RNA Extraction was done from control and treated yeast samples. The quantification of the RNA check by Nano-Drop, which was used further for C-DNA preparation. The online NCBI Primers-BLAST was used to design the primers of LDs recognizing genes (SEI1, LDH1, and PLN1) for qRT-PCR. The real-time RCR based primer of all suggested proteins designed, as shown in Table 2. The standardization of Real-Time PCR was done on Roche’s LightCycler® 480 using SYBR green dye. The primers used under prescribed cycling conditions (X. Wang et al. 2020) and the melt curve analysis at 95 °C for 10 sec, 60 °C for 1 min or 5 °C below lowest primer TM, and 4 °C for hold. Analyses of the data and plotting of these curves with software LightCycler® 480 SW 1.5. Data Analysis. For the real-time data analysis, assume parasitemia calculated using 2(-ΔΔCt) with the help of LightCycler® 480 SW 1.5 and Microsoft Office Excel 2007.

**Table 2.**
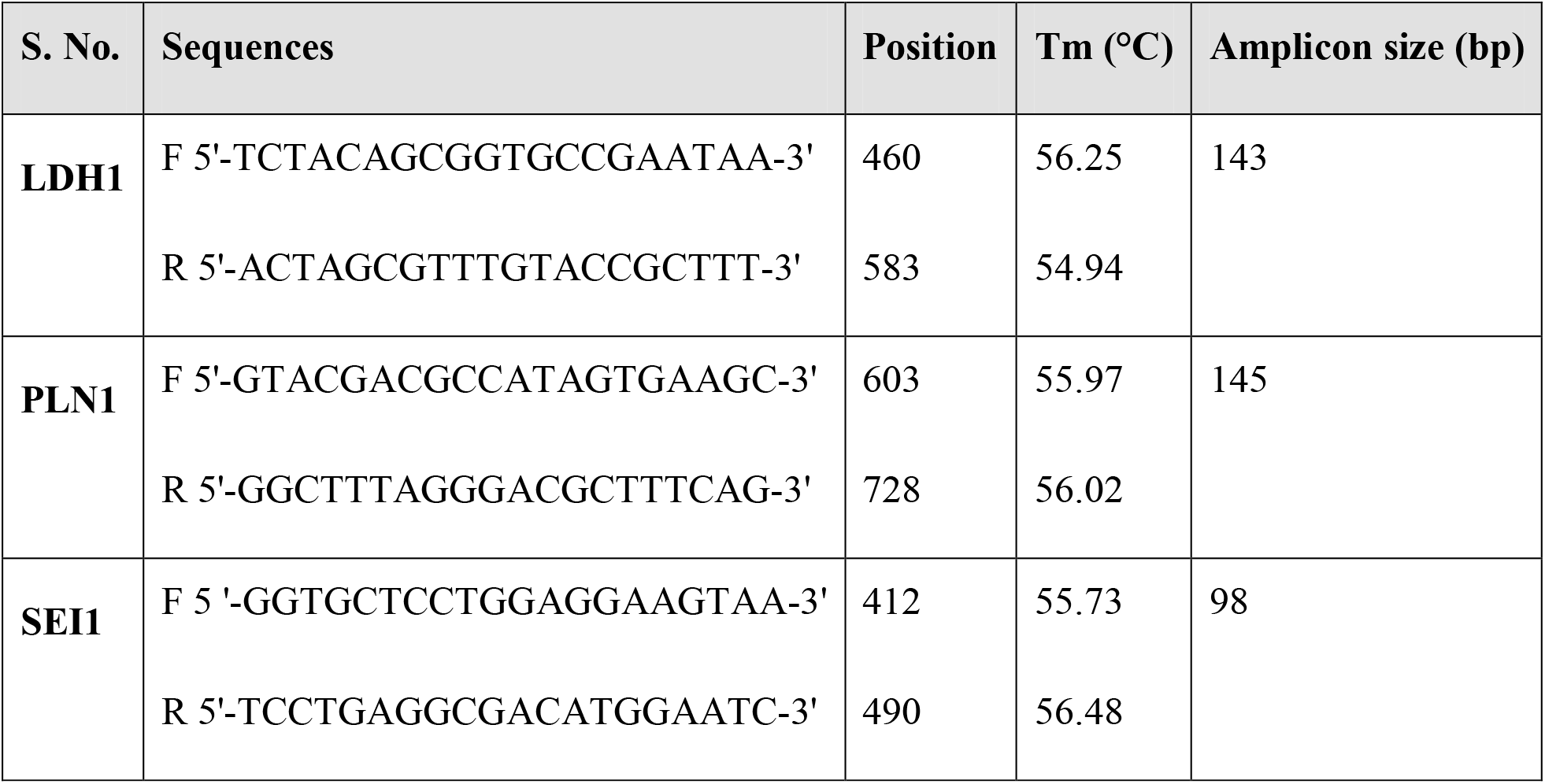
Lipid Droplet specific Real-time PCR primers

### Spot assay

The yeast cells under control and treated samples suspend in the sterile water under fivefold serial dilutions of each yeast culture. Afterward, 2 μL of each cell suspension was inoculated on a solid YPD medium and incubate at 37°C. Colony growth differences review after 72 h incubation.

### Confocal microscopy

MitoTracker Deep Red purchased from Thermo Fisher containing identical 20 vials. Each vial (50 μg) of MitoTracker Deep Red dissolved with 91.98 μl of high-quality DMSO to make a 1mM stock solution. Mitochondrial assessment can obtain by diluting the stock solution to 500nM as prescribed. Cells under control and treated conditions were washed with PBS 2 times and re-suspended in 100 μl of PBS solution. The sub-stock solution of MitoTracker Deep Red was added to both samples and incubated for 30 min at 37°C in the dark. Incubated samples were further washed with PBS and fixed in 4% paraformaldehyde. The Cell Navigator Fluorimetric Lipid Droplet assay kit purchase from AAT Bioquest’s, a robust tool that quantitatively measures the cell’s LDs accumulation (Greenspan and Fowler 1985). The Nile red is a lipophilic strain that detects primarily intracellular LDs (Greenspan, Mayer, and Fowler 1985) and increases robustness. Stained cells under every condition mounted onto a glass slide that contains 16 mm glass coverslips. After complete adhesion, the cells were visualized by Nikon A1R inverted confocal microscope (Nikon Corp, Japan). Localization of mitochondria and LDs in the cellular environment was analyzed using 550 nm and 644 nm laser. Cells under confocal microscopy were selected based on the shape and characteristic features for identification. The stained cells were analyzed using NIS elements software.

### Flow Cytometry

Cells under control and treated conditions were centrifuged, re-suspended in PBS, and studied by forward scatter (FSC) versus side scatter (SSC). The MitoTracker deep red probe was used for mitochondria bioprocess analysis using excitation wavelength at 644 nm and emission at 665 nm. The lipid droplet analysis was done using Nile red as a probe using 550 nm excitation and 640 nm emission. Besides, the intracellular ROS level was analyzed using non-fluorescent dihydrorhodamine 123 (DHR 123) dye, which converts to fluorescent rhodamine 123 under oxidative stress (Pérez-Gallardo et al. 2013). CFSE (5[6]-Carboxyfluorescein Diacetate Succinimidyl Ester) cell proliferation kit purchased from Bio-Rad. CFSE cell proliferation kit was excited on 492 nm, and emission was calculated on 517 nm (Quah, Warren, and Parish 2007). The yeast cells under control and treated samples were taken from the 48th hour of the experiment. The cells were harvested, washed, and eventually re-suspended in PBS buffer followed by incubation in the dark.

## Graphical Abstract

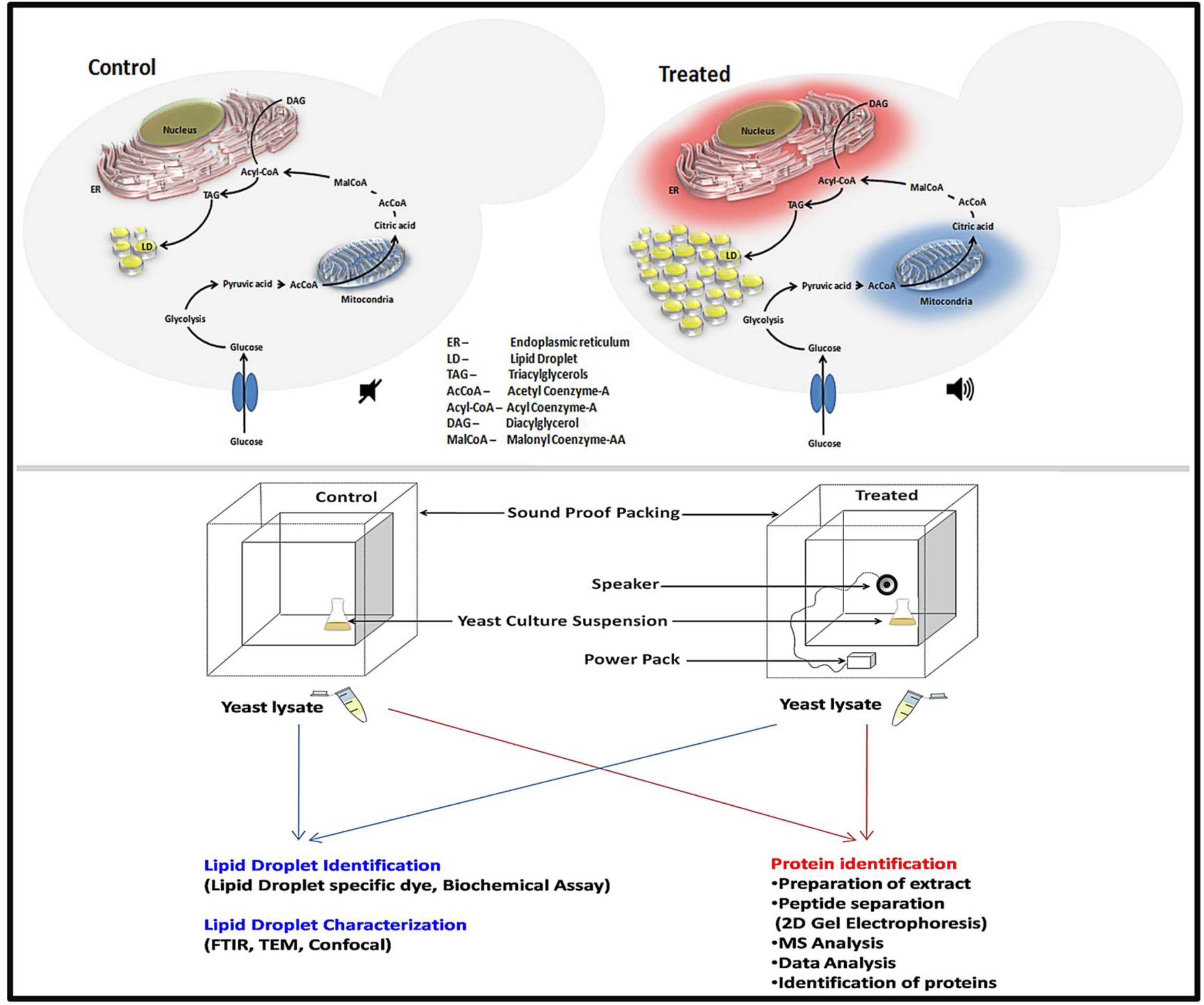

## Supplementary Materials legends

**Supplementary Audio SA1.** The audio file enchanting “AUM” as inducer. The lower and higher intensity adjust by Audacity software (www.audacityteam.org)

**Supplementary Figure SF1.** Three-dimensional representation of 2D gel image as shown in control and treated sample by Melanie software. Spots were selected precisely using this software for further analysis of MALDI experiments.

**Supplementary Figure SF2.** The WikiPathways pathway diagram for Glycolysis and gluconeogenesis pathway, which merge later in the mitochondrion. The noise expressed protein is used as input for pathway generation.

**Supplementary Table ST1.** Protein-protein interaction among noise expressed protein: STRING Analysis

**Supplementary Table ST2.** MCODE suggested cluster among expressed proteins using STRING Protein-protein interaction statistics.

**Supplementary Table ST3.** Gene Ontogeny and KEGG pathway analysis of noise expressed proteins.

## References

Abu-Farha, Mohamed et al. 2020. “The Role of Lipid Metabolism in COVID-19 Virus Infection and as a Drug Target.” International Journal of Molecular Sciences 21(10): 3544.

Aggio, Raphael Bastos Mereschi, Victor Obolonkin, and Silas Granato Villas-Bôas. 2012. “Sonic Vibration Affects the Metabolism of Yeast Cells Growing in Liquid Culture: A Metabolomic Study.” Metabolomics 8(4): 670–78.

Algers, B., I. Ekesbo, and S. Strömberg. 1978. “The Impact of Continuous Noise on Animal Health.” Acta Veterinaria Scandinavica. Supplementum (68): 1–26.

Alptekin, Ertan, Mustafa Canakci, and Huseyin Sanli. 2014. “Biodiesel Production from Vegetable Oil and Waste Animal Fats in a Pilot Plant.” Waste Management 34(11): 2146–54.

Appel, R. D. et al. 1997. “Melanie II--a Third-Generation Software Package for Analysis of Two-Dimensional Electrophoresis Images: I. Features and User Interface.” Electrophoresis 18(15): 2724–34.

Aro, Eva-Mari. 2016. “From First Generation Biofuels to Advanced Solar Biofuels.” Ambio 45(S1): 24–31.

Atadashi, I. M., M. K. Aroua, A. R. Abdul Aziz, and N. M. N. Sulaiman. 2012. “The Effects of Water on Biodiesel Production and Refining Technologies: A Review.” Renewable and Sustainable Energy Reviews 16(5): 3456–70.

Bader, Gary D., and Christopher W. V. Hogue. 2003. “An Automated Method for Finding Molecular Complexes in Large Protein Interaction Networks.” BMC bioinformatics 4: 2.

Bailey, Andrew P. et al. 2015. “Antioxidant Role for Lipid Droplets in a Stem Cell Niche of Drosophila.” Cell 163(2): 340–53.

Bartz, René et al. 2007. “Lipidomics Reveals That Adiposomes Store Ether Lipids and Mediate Phospholipid Traffic.” Journal of Lipid Research 48(4): 837–47.

Beopoulos, Athanasios, and Jean-Marc Nicaud. 2012. “Yeast: A New Oil Producer?” Oléagineux, Corps gras, Lipides 19(1): 22–28.

Bersuker, Kirill, and James A. Olzmann. 2017. “Establishing the Lipid Droplet Proteome: Mechanisms of Lipid Droplet Protein Targeting and Degradation.” Biochimica Et Biophysica Acta. Molecular and Cell Biology of Lipids 1862(10 Pt B): 1166–77.

Berth, Matthias, Frank Michael Moser, Markus Kolbe, and Jörg Bernhardt. 2007. “The State of the Art in the Analysis of Two-Dimensional Gel Electrophoresis Images.” Applied Microbiology and Biotechnology 76(6): 1223–43.

Bhuiya, M. M. K. et al. 2014. “Second Generation Biodiesel: Potential Alternative to - Edible Oil-Derived Biodiesel.” Energy Procedia 61: 1969–72.

Blake, William J., Mads KAErn, Charles R. Cantor, and J. J. Collins. 2003. “Noise in Eukaryotic Gene Expression.” Nature 422(6932): 633–37.

Bosch, Marta et al. 2020. “Mammalian Lipid Droplets Are Innate Immune Hubs Integrating Cell Metabolism and Host Defense.” Science 370(6514): eaay8085.

Carere, Carlo R., Richard Sparling, Nazim Cicek, and David B. Levin. 2008. “Third Generation Biofuels via Direct Cellulose Fermentation.” International Journal of Molecular Sciences 9(7): 1342–60.

Castelhano-Carlos, M. J., and V. Baumans. 2009. “The Impact of Light, Noise, Cage Cleaning and in-House Transport on Welfare and Stress of Laboratory Rats.” Laboratory Animals 43(4): 311–27.

Cermelli, Silvia, Yi Guo, Steven P. Gross, and Michael A. Welte. 2006. “The Lipid - Droplet Proteome Reveals That Droplets Are a Protein-Storage Depot.” Current biology: CB 16(18): 1783–95.

Chen, Xiaoyi et al. 2015. “Automated Flow Cytometric Analysis across Large Numbers of Samples and Cell Types.” Clinical Immunology (Orlando, Fla.) 157(2): 249–60.

Cheung, Winsome et al. 2010. “Rotaviruses Associate with Cellular Lipid Droplet Components to Replicate in Viroplasms, and Compounds Disrupting or Blocking Lipid Droplets Inhibit Viroplasm Formation and Viral Replication.” Journal of Virology 84(13): 6782–98.

Coffey, Caroline M. et al. 2006. “Reovirus Outer Capsid Protein Micro1 Induces Apoptosis and Associates with Lipid Droplets, Endoplasmic Reticulum, and Mitochondria.” Journal of Virology 80(17): 8422–38.

Colpitts, Che C., Joachim Lupberger, Christian Doerig, and Thomas F. Baumert. 2015. “Host Cell Kinases and the Hepatitis C Virus Life Cycle.” Biochimica Et Biophysica Acta 1854(10 Pt B): 1657–62.

Dahlqvist, A. et al. 2000. “Phospholipid:Diacylglycerol Acyltransferase: An Enzyme That Catalyzes the Acyl-CoA-Independent Formation of Triacylglycerol in Yeast and Plants.” Proceedings of the National Academy of Sciences of the United States of America 97(12): 6487–92.

Debelyy, Mykhaylo O. et al. 2011. “Involvement of the Saccharomyces Cerevisiae Hydrolase Ldh1p in Lipid Homeostasis.” Eukaryotic Cell 10(6): 776–81.

Di Talia, Stefano et al. 2007. “The Effects of Molecular Noise and Size Control on Variability in the Budding Yeast Cell Cycle.” Nature 448(7156): 947–51.

Eling, T. E., and W. C. Glasgow. 1994. “Cellular Proliferation and Lipid Metabolism: Importance of Lipoxygenases in Modulating Epidermal Growth Factor-Dependent Mitogenesis.” Cancer Metastasis Reviews 13(3–4): 397–410.

Faber, B. C. et al. 2001. “Identification of Genes Potentially Involved in Rupture of Human Atherosclerotic Plaques.” Circulation Research 89(6): 547–54.

Farese, Robert V., and Tobias C. Walther. 2009. “Lipid Droplets Finally Get a Little R-E-S-P-E-C-T.” Cell 139(5): 855–60.

Fecchi, Katia et al. 2020. “Coronavirus Interplay With Lipid Rafts and Autophagy Unveils Promising Therapeutic Targets.” Frontiers in Microbiology 11: 1821.

Filipe, Ana, and John McLauchlan. 2015. “Hepatitis C Virus and Lipid Droplets: Finding a Niche.” Trends in Molecular Medicine 21(1): 34–42.

Forfang, Kristin et al. 2017. “FTIR Spectroscopy for Evaluation and Monitoring of Lipid Extraction Efficiency for Oleaginous Fungi.” PloS One 12(1): e0170611.

Fraser, Hunter B. et al. 2004. “Noise Minimization in Eukaryotic Gene Expression.” PLOS Biology 2(6): e137.

Fujimoto, Toyoshi, and Robert G. Parton. 2011. “Not Just Fat: The Structure and Function of the Lipid Droplet.” Cold Spring Harbor Perspectives in Biology 3(3).

Gao, Qiang et al. 2017. “Pet10p Is a Yeast Perilipin That Stabilizes Lipid Droplets and Promotes Their Assembly.” Journal of Cell Biology 216(10): 3199–3217.

Ghosh, Ritesh et al. 2016. “Exposure to Sound Vibrations Lead to Transcriptomic, Proteomic and Hormonal Changes in Arabidopsis.” Scientific Reports 6: 33370.

Goodman, Joel M. 2008. “The Gregarious Lipid Droplet.” The Journal of Biological Chemistry 283(42): 28005–9.

Greenspan, P., and S. D. Fowler. 1985. “Spectrofluorometric Studies of the Lipid Probe, Nile Red.” Journal of Lipid Research 26(7): 781–89.

Greenspan, P., E. P. Mayer, and S. D. Fowler. 1985. “Nile Red: A Selective Fluorescent Stain for Intracellular Lipid Droplets.” The Journal of Cell Biology 100(3): 965–73.

Hartley, Andrew M. et al. 2019. “Structure of Yeast Cytochrome c Oxidase in a Supercomplex with Cytochrome Bc 1.” Nature Structural & Molecular Biology 26(1): 78–83.

Herker, Eva, and Melanie Ott. 2012. “Emerging Role of Lipid Droplets in Host/Pathogen Interactions.” The Journal of Biological Chemistry 287(4): 2280–87.

Hooshangi, Sara, Stephan Thiberge, and Ron Weiss. 2005. “Ultrasensitivity and Noise Propagation in a Synthetic Transcriptional Cascade.” Proceedings of the National Academy of Sciences of the United States of America 102(10): 3581–86.

Kight, Caitlin R., and John P. Swaddle. 2011. “How and Why Environmental Noise Impacts Animals: An Integrative, Mechanistic Review.” Ecology Letters 14(10): 1052–61.

Koll, Hans et al. 1992. “Antifolding Activity of Hsp60 Couples Protein Import into the Mitochondrial Matrix with Export to the Intermembrane Space.” Cell 68(6): 1163–75.

Kushnirov, V. V. 2000. “Rapid and Reliable Protein Extraction from Yeast.” Yeast (Chichester, England) 16(9): 857–60.

Kutmon, Martina, Samad Lotia, Chris T. Evelo, and Alexander R. Pico. 2014. “WikiPathways App for Cytoscape: Making Biological Pathways Amenable to Network Analysis and Visualization.” F1000Research 3: 152.

Leber, R. et al. 1998. “Dual Localization of Squalene Epoxidase, Erg1p, in Yeast Reflects a Relationship between the Endoplasmic Reticulum and Lipid Particles.” Molecular Biology of the Cell 9(2): 375–86.

Lehner, Ben. 2008. “Selection to Minimise Noise in Living Systems and Its Implications for the Evolution of Gene Expression.” Molecular Systems Biology 4: 170.

Li, Zhihuan et al. 2012. “Lipid Droplets Control the Maternal Histone Supply of Drosophila Embryos.” Current biology: CB 22(22): 2104–13.

Lin, Hui et al. 2013. “Genetic Engineering of Microorganisms for Biodiesel Production.” Bioengineered 4(5): 292–304.

Liu, Jian, Jean-Marie François, and Jean-Pascal Capp. 2016. “Use of Noise in Gene Expression as an Experimental Parameter to Test Phenotypic Effects.” Yeast (Chichester, England) 33(6): 209–16.

Lü, Jing, Con Sheahan, and Pengcheng Fu. 2011. “Metabolic Engineering of Algae for Fourth Generation Biofuels Production.” Energy & Environmental Science 4(7): 2451–66.

Lyn, Rodney K. et al. 2013. “Bidirectional Lipid Droplet Velocities Are Controlled by Differential Binding Strengths of HCV Core DII Protein.” PLoS ONE 8(11). https://www.ncbi.nlm.nih.gov/pmc/articles/PMC3815211/ (November 16, 2020).

Meneghetti, Simoni M. Plentz et al. 2007. “Biodiesel Production from Vegetable Oil Mixtures: Cottonseed, Soybean, and Castor Oils.” Energy & Fuels 21(6): 3746–47.

Mihoubi, Wafa, Emna Sahli, Ali Gargouri, and Caroline Amiel. 2017. “FTIR Spectroscopy of Whole Cells for the Monitoring of Yeast Apoptosis Mediated by P53 Over-Expression and Its Suppression by Nigella Sativa Extracts.” PloS One 12(7): e0180680.

Miyanari, Yusuke et al. 2007. “The Lipid Droplet Is an Important Organelle for Hepatitis C Virus Production.” Nature Cell Biology 9(9): 1089–97.

Mundt, Max, Alexander Anders, Seán M. Murray, and Victor Sourjik. 2018. “A System for Gene Expression Noise Control in Yeast.” ACS synthetic biology 7(11): 2618–26.

Murphy, Samantha, Sally Martin, and Robert G. Parton. 2009. “Lipid Droplet-Organelle Interactions; Sharing the Fats.” Biochimica Et Biophysica Acta 1791(6): 441–47.

Nachman, Iftach, Aviv Regev, and Sharad Ramanathan. 2007. “Dissecting Timing Variability in Yeast Meiosis.” Cell 131(3): 544–56.

Nardacci, Roberta et al. 2020. SARS-CoV-2 Cytopathogenesis in Cultured Cells and in COVID-19 Autoptic Lung, Evidences of Lipid Involvement. In Review. preprint. https://www.researchsquare.com/article/rs-39274/v1 (November 16, 2020).

Newman, John R. S. et al. 2006. “Single-Cell Proteomic Analysis of S. Cerevisiae Reveals the Architecture of Biological Noise.” Nature 441(7095): 840–46.

Niehus, Xochitl, Leticia Casas-Godoy, Francisco J. Rodríguez-Valadez, and Georgina Sandoval. 2018. “Evaluation of Yarrowia Lipolytica Oil for Biodiesel Production: Land Use Oil Yield, Carbon, and Energy Balance.” Journal of Lipids 2018: 6393749.

Olofsson, Sven-Olof et al. 2011. “The Formation of Lipid Droplets: Possible Role in the Development of Insulin Resistance/Type 2 Diabetes.” Prostaglandins, Leukotrienes, and Essential Fatty Acids 85(5): 215–18.

Paul, Rajib et al. 2017. “Cholesterol Contributes to Dopamine-Neuronal Loss in MPTP Mouse Model of Parkinson’s Disease: Involvement of Mitochondrial Dysfunctions and Oxidative Stress.” PloS One 12(2): e0171285.

Pereira-Dutra, Filipe S., Livia Teixeira, Maria Fernanda de Souza Costa, and Patrícia T. Bozza. 2019. “Fat, Fight, and beyond: The Multiple Roles of Lipid Droplets in Infections and Inflammation.” Journal of Leukocyte Biology 106(3): 563–80.

Pérez-Gallardo, Rocio V. et al. 2013. “Reactive Oxygen Species Production Induced by Ethanol in Saccharomyces Cerevisiae Increases Because of a Dysfunctional Mitochondrial Iron-Sulfur Cluster Assembly System.” FEMS yeast research 13(8): 804–19.

Quah, Ben J. C., Hilary S. Warren, and Christopher R. Parish. 2007. “Monitoring Lymphocyte Proliferation in Vitro and in Vivo with the Intracellular Fluorescent Dye Carboxyfluorescein Diacetate Succinimidyl Ester.” Nature Protocols 2(9): 2049–56.

Radulovic, Maja et al. 2013. “The Emergence of Lipid Droplets in Yeast: Current Status and Experimental Approaches.” Current Genetics 59(4): 231–42.

Ranford, Julia C., Anthony R.M. Coates, and Brian Henderson. 2000. “Chaperonins Are Cell-Signalling Proteins: The Unfolding Biology of Molecular Chaperones.” Expert Reviews in Molecular Medicine 2(8): 1–17.

Samsa, Marcelo M. et al. 2009. “Dengue Virus Capsid Protein Usurps Lipid Droplets for Viral Particle Formation.” PLoS pathogens 5(10): e1000632.

Sarvaiya, Niral, and Vijay Kothari. 2015. “Effect of Audible Sound in Form of Music on Microbial Growth and Production of Certain Important Metabolites.” Microbiology 84(2): 227–35.

Shaobin, Gu et al. 2010. “A Pilot Study of the Effect of Audible Sound on the Growth of Escherichia Coli.” Colloids and Surfaces. B, Biointerfaces 78(2): 367–71.

da Silva Gomes Dias, Suelen et al. 2020. Lipid Droplets Fuel SARS-CoV-2 Replication and Production of Inflammatory Mediators. Immunology. preprint. http://biorxiv.org/lookup/doi/10.1101/2020.08.22.262733 (November 16, 2020).

Singaram, Lakshmanan. 2009. “Biodiesel: An Eco-Friendly Alternate Fuel for the Future: A Review.” Thermal Science 13(3): 185–99.

Steen, Eric J. et al. 2010. “Microbial Production of Fatty-Acid-Derived Fuels and Chemicals from Plant Biomass.” Nature 463(7280): 559–62.

Stobart, A. K., S. Stymne, and S. Höglund. 1986. “Safflower Microsomes Catalyse Oil Accumulation in Vitro: A Model System.” Planta 169(1): 33–37.

Sui, Xuewu et al. 2018. “Cryo-Electron Microscopy Structure of the Lipid Droplet-Formation Protein Seipin.” The Journal of Cell Biology 217(12): 4080–91.

Szklarczyk, Damian et al. 2015. “STRING V10: Protein-Protein Interaction Networks, Integrated over the Tree of Life.” Nucleic Acids Research 43(Database issue): D447–452.

Sztalryd, Carole, and Dawn L. Brasaemle. 2017. “The Perilipin Family of Lipid Droplet Proteins: Gatekeepers of Intracellular Lipolysis.” Biochimica et biophysica acta 1862(10 Pt B): 1221–32.

Tai, Mitchell, and Gregory Stephanopoulos. 2013. “Engineering the Push and Pull of Lipid Biosynthesis in Oleaginous Yeast Yarrowia Lipolytica for Biofuel Production.” Metabolic Engineering 15: 1–9.

Tatsumi, Takayuki et al. 2018. “Forced Lipophagy Reveals That Lipid Droplets Are Required for Early Embryonic Development in Mouse.” Development (Cambridge, England) 145(4).

Thoms, Sven et al. 2011. “The Putative Saccharomyces Cerevisiae Hydrolase Ldh1p Is Localized to Lipid Droplets.” Eukaryotic Cell 10(6): 770–75.

Tkacik, Gasper, Curtis G. Callan, and William Bialek. 2008. “Information Flow and Optimization in Transcriptional Regulation.” Proceedings of the National Academy of Sciences of the United States of America 105(34): 12265–70.

Tsai, Ching-Sung, Suryang Kwak, Timothy L. Turner, and Yong-Su Jin. 2015. “Yeast Synthetic Biology Toolbox and Applications for Biofuel Production.” FEMS yeast research 15(1): 1–15.

Veltman, Douwe M., and Robert H. Insall. 2010. “WASP Family Proteins: Their Evolution and Its Physiological Implications.” Molecular Biology of the Cell 21(16): 2880–93.

Villareal, Valerie A., Mary A. Rodgers, Deirdre A. Costello, and Priscilla L. Yang. 2015. “Targeting Host Lipid Synthesis and Metabolism to Inhibit Dengue and Hepatitis C Viruses.” Antiviral Research 124: 110–21.

Wang, Huajin et al. 2016. “Seipin Is Required for Converting Nascent to Mature Lipid Droplets.” eLife 5.

Wang, Xuefei, Dean A. Glawe, David M. Weller, and Patricia A. Okubara. 2020. “Real-Time PCR Assays for the Quantification of Native Yeast DNA in Grape Berry and Fermentation Extracts.” Journal of Microbiological Methods 168: 105794.

Welte, Michael A. 2015. “Expanding Roles for Lipid Droplets.” Current biology: CB 25(11): R470–481.

Yan, J. X. et al. 2000. “A Modified Silver Staining Protocol for Visualization of Proteins Compatible with Matrix-Assisted Laser Desorption/Ionization and Electrospray Ionization-Mass Spectrometry.” Electrophoresis 21(17): 3666–72.

Yang, Po-Lin, Tzu-Han Hsu, Chao-Wen Wang, and Rey-Huei Chen. 2016. “Lipid Droplets Maintain Lipid Homeostasis during Anaphase for Efficient Cell Separation in Budding Yeast” ed. Anne Spang. Molecular Biology of the Cell 27(15): 2368–80.

Zoni, Valeria et al. 2019. “Lipid Droplet Biogenesis Is a Liquid Phase Separation Spatially Regulated by Seipin and Membrane Curvature.” bioRxiv: 777466

